# Bidirectional relationship between attentional deficits and escalation of nicotine intake

**DOI:** 10.1101/2023.12.22.572989

**Authors:** Caroline Vouillac-Mendoza, Serge H. Ahmed, Karine Guillem

**Author notes:** **Corresponding author**: Karine Guillem, PhD Université de Bordeaux Institut de Neurosciences Cognitives et Intégratives d’Aquitaine, CNRS UMR 5287 Bâtiment BBS - 2ème étage 2, Rue du Dr Hoffmann Martinot 33000 Bordeaux, France.

## Abstract

Smoking addicts have deficits in cognition, in particular deficits in attention, even long after smoking cessation. It is not clear however, whether deficits are a cause or a consequence, or both, of chronic nicotine use. Here we set out a series of experiments in rats to address this question and, more specifically, to assess the long-term effects of exposure to and withdrawal from chronic nicotine self-administration on attentional performance. Animals were trained in a 5-choice serial reaction time task to probe individual attentional performance and, then, were given access to a fixed versus increasing dose of intravenous nicotine for self-administration, a differential dose procedure known to induce two between-session patterns of nicotine intake: a stable versus escalation pattern. Attentional performance was measured daily before, during and also 24-h after chronic access to the differential dose procedure of nicotine self-administration. We found that pre-existing individual variation in attentional performance predicts individual vulnerability to develop escalation of nicotine intake. Moreover, while chronic nicotine self-administration increases attention, withdrawal from nicotine intake escalation induces attentional deficits, a withdrawal effect that is dose-dependently reversed by acute nicotine. Together, these results suggest that pre-existing individual variation in attentional performance predicts individual vulnerability to develop escalation of nicotine intake, and that part of the motivation for using nicotine during escalation might be to alleviate withdrawal-induced attentional deficits.

## Introduction

Beside its well-established reinforcing effects, nicotine also acts as a cognitive enhancer in humans and animals [1,2], a psychoactive effect that contributes to both the initiation and maintenance of tobacco use. In contrast, heavy cigarette smokers have deficits in cognitive function, in particular deficits in attention during withdrawal that can persist even long after smoking cessation [3–6]. Whether these attentional deficits are a cause or a consequence, or both, of chronic nicotine use remain to be elucidated. In theory, these attentional deficits could either pre-exist to nicotine use thereby explaining smokers’ poor attentional performance, or they could directly result from chronic nicotine use. Alternatively, and more likely, these two phenomena may causally influence each other through a spiraling process. That is, future smokers may have some minor pre-existing attentional deficits which predispose them to start smoking, and afterwards nicotine use in these individuals will worsen these attentional deficits thereby further increasing the need of using nicotine to enhance cognition.

In rats, attentional functions have been extensively studied using the 5-choice serial reaction time task (5-CSRTT) [7,8]. To date, most of the rat’s studies using the 5-CSRTT task have focused on the short-term effects of acute nicotine exposure and have consistently shown that nicotine improves sustained attention [1,9–12]. Comparatively fewer studies have investigated the long-term effects of chronic nicotine exposure and withdrawal on 5-CSRTT attention performance and have shown that the enhancing effect of nicotine on attention remains during chronic use while withdrawal from chronic nicotine use induces attentional deficits [11,13,14]. However, these withdrawal deficits were relatively mild and lasted only few hours [15] which contrasts with the enduring attentional dysfunctions seen in abstinent smokers. One factor that may explain this difference could be the procedure used in rats for chronic nicotine exposure (i.e., experimenter-delivered injections by osmotic mini-pumps) that may not be sufficiently efficient in producing some key addiction-related behavioral changes, such as, escalation of drug self-administration [16–18]. Interestingly, we have recently developed a robust rat model of escalation of nicotine intake and motivation by giving animals access to a progressively increasing dose of nicotine for self-administration [19].

The aim of the present study is to use this model to assess: i) how individual variation in attentional performance can relate to individual vulnerability to nicotine intake escalation; ii) whether and to what extent withdrawal from nicotine intake escalation can induce attentional deficits; and, finally, iii) whether these deficits can be reversed by returning to nicotine use. Animals were initially trained in the 5-CSRTT under a variable stimulus duration procedure (variable SD) to probe individual attentional performance. Then, they had access to either a fixed unit dose of nicotine for self-administration (i.e., 30 µg/kg/injection; no-escalation group = NoES) or to an increasing unit dose of nicotine for self-administration (i.e., from 30 μg to 240 μg/kg/injection; escalation group = ES) to precipitate drug intake escalation. Overall, our data suggest that pre-existing individual variation in attentional performance predicts individual vulnerability to develop escalation of nicotine intake, and that part of the motivation for using nicotine during escalation might be to alleviate withdrawal-induced attentional deficits.

## Material and Methods

### Subjects

A total of 68 adult male Wistar rats (300-325 g at the beginning of experiments, Charles River, Lyon, France) were used. Rats were housed in groups of 2 and were maintained in a light-(reverse light-dark cycle), humidity-(60 ± 20%) and temperature-controlled vivarium (21 ± 2°C), with water available ad libitum. Animals were food restricted (∼ 12-16 g/rat) to maintain at least 95% of their free feeding body weight and food ration were given 3 hours after the end of the session. All behavioral testing occurred during the dark phase of the light-dark cycle. Home cages were enriched with a nylon gnawing bone and a cardboard tunnel (Plexx BV, The Netherlands). All experiments were carried out in accordance with institutional and international standards of care and use of laboratory animals [UK Animals (Scientific Procedures) Act, 1986; and associated guidelines; the European Communities Council Directive (2010/63/UE, 22 September 2010) and the French Directives concerning the use of laboratory animals (décret 2013-118, 1 February 2013)]. The animal facility has been approved by the Committee of the Veterinary Services Gironde, agreement number B33-063-922.

### Five-choice serial reaction time task (5-CSRTT)

#### Apparatus

Height identical five-hole nose poke operant chambers (30 x 40 x 36 cm) housed in sound-insulating and ventilated cubicles were used for 5-CSRTT testing and training (Imétronic, Pessac, France) (Fig. 1), as described previously [20]. Each chamber was equipped with stainless steel grid floors and a white house light mounted in the center of the roof. One side wall was curved inward to present an array of five circular holes (2.5 cm sides, 4 cm deep and positioned 2 cm above the grid floor), each with an internal light-emitting diode and an infrared sensor for detecting nose insertion. The opposite side wall was equipped with a delivery port with a drinking cup mounted on the midline. The delivery port was illuminated with a white light diode mounted 8.5 cm above the drinking cup. Each chamber was also equipped with a syringe pump placed outside on the top of the cubicle and connected to the drinking cup via a silastic tubing (Dow Corning Corporation, Michigan, USA).

**Figure 1.**
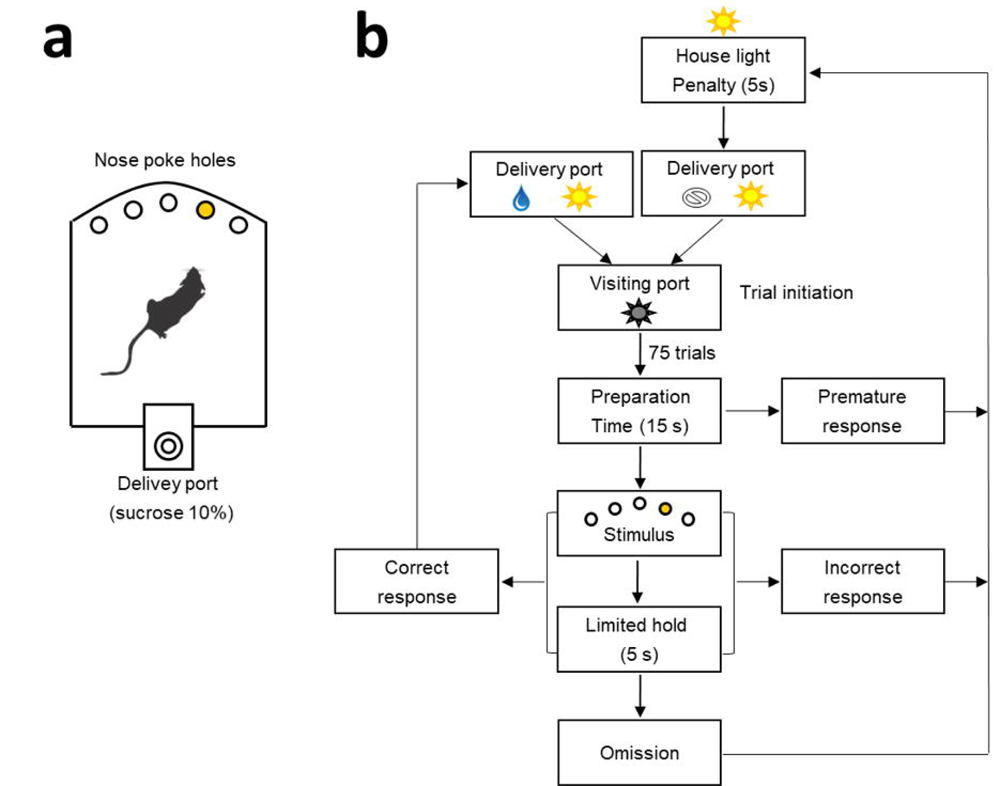
5-CSRTT chamber and general procedure. (**a**) Schematic of the 5-CSRTT apparatus and (**b**) diagram showing the sequence of events during a 5-CSRTT training session. To initiate a trial, rats had to turn off the illuminated delivery port by visiting it (i.e., exit after entering). After a fixed preparation time of 15 sec, a brief stimulus is presented in one of the 5 holes on the opposite curved wall. Animals had to nose-poke the illuminated hole (correct response), either during the stimulus presentation or within a post-stimulus limited hold (LH) period of 5 seconds, to receive a sucrose reward (0.1 ml of 10% sucrose). If they respond to the wrong hole (incorrect response), respond before the stimulus is presented (premature response), or fail to respond (omission), they are punished by a time-out penalty of 5 seconds signaled by turning on the house light. After that, the delivery port is switched on and rats have to visit it (i.e., exit after entering) to initiate the next trial.

## 5-CSRTT training and testing

Animals were trained with a maximum number of 75 consecutive self-paced trials per session or a maximum duration of 60 min whichever came first, as previously described [20] (Fig. 1a, b). Briefly, the opportunity to self-initiate a trial was signaled by turning on the white light diode in the delivery port and rats had to turn off the illuminated port by visiting it (i.e., exit after entering) within one minute to initiate a trial. If no visit occurred within the imparted time, the port was turned off automatically and this marked trial onset. After a fixed preparation time of 15 sec, a brief light stimulus was presented behind one of the 5 holes on the opposite curved wall (pseudorandom selection across trials). To receive a reward (0.1 ml of 10% sucrose), animals had to nose-poke the illuminated hole either during the stimulus presentation or within a post-stimulus limited hold (LH) period of 5 seconds (Fig. 1b). Correct nose-poke responses immediately turned off the light stimulus (if still on), turned back on the light in the delivery port and triggered the delivery of sucrose into the drinking cup. Incorrect nose-poke responses in one of the dark holes were not rewarded by sucrose, but were punished by a time-out penalty of 5 seconds signaled by turning on the house light. If animals failed to respond in any of the holes during a trial, this was considered an omission response. Omissions, like incorrect responses, were punished by a signaled 5-s time-out penalty. At the end of the time-out penalty, the house light was switched off and the light in the delivery port was turned back on for the next trial. During training, the duration of light stimulus was initially set to 30 s and progressively decreased across sessions to 1 s until the subject met performance criteria (omissions < 30%; accuracy > 60%; number of self-initiated trials > 50).

Once performance was stable with a light stimulus of 1 s, attentional demand was further increased within-session by decreasing the duration of the light stimulus in the following order: 1, 0.75, and 0.5 seconds. Each stimulus duration (SD) was tested during at least 25 trials per session. Task performance was reflected in the following behavioral measures: (1) % accuracy ([100 x correct responses] / [correct responses + incorrect responses]); % omission (100 x number of omissions / total self-initiated trials); premature responses (number of responses that occurred before the presentation of the light stimulus); latency of correct and incorrect responses; and, finally, reward latency (i.e., latency between correct responses and contact with the drinking cup). Animals were tested in this SD procedure during several daily sessions until stabilization of performance. Attentional performances were measured at different times depending on the experiments (see Specific experiments).

### Differential dose procedure of intravenous nicotine self-administration

Fourteen identical operant chambers (30 x 40 x 36 cm) were used for nicotine self-administration testing and training (Imétronic, Pessac, France). Each chamber was equipped with two retractable metal levers on opposite panels of the chamber and a corresponding white cue light above each lever. Animals were surgically prepared with an indwelling silastic catheter (Dow Corning Corporation, Michigan, USA) in the right jugular vein under deep anesthesia (mixture of ketamine 100 mg/kg and xylazine 15 mg/kg, i.p). After surgery, animals were flushed daily with 0.2 ml of an ampicillin solution (0.1 g/ml) containing heparin (300 IU/ml) to maintain patency. Then, animals were trained to self-administer nicotine for 15 days (30 µg/ injection; 40 µl in 1s; fixed-ratio 1 schedule of reinforcement; 3-h sessions). All self-administration sessions began with extension of the operant lever and ended with its retraction after 3 h. Intravenous delivery of nicotine began immediately after completion of the lever press and was accompanied by illumination of the light cue above the lever for 20 s. Responses during the light cue were recorded but had no programmed consequence. Then, animals were divided in two groups and subjected to a differential dose procedure of nicotine self-administration that was previously shown to induce two between-session patterns of nicotine intake: a no-escalation versus escalation pattern [19]. Briefly, during this differential dose procedure, the unit dose of nicotine remained at the initial dose of 30 µg/injection for 30 additional sessions in one group of rats (i.e., no-escalation group = No ES,), while it progressively increased to 120 µg and then to 240 µg/injection for 15 sessions at each dose in the other group (i.e., escalation group = ES) (Fig. 3a). This increase in the unit dose of nicotine available was done by increasing the injection volume (to 160 and 320 µl).

### Specific experiments

#### Experiment 1: relationship between individual attentional performance and individual vulnerability to escalation of nicotine intake

To study this relationship, animals (*n* = 41; No ES: *n* = 12; ES: *n* = 29) were initially tested in the 5-CSRTT under the SD procedure, as described above, before being given access to chronic nicotine for self-administration under the differential dose procedure.

#### Experiment 2: effects of chronic nicotine self-administration and withdrawal on attentional performance

Experiment 2 was conducted in a separate group of rats (*n* = 27; NoES: *n* = 12 and ES: *n*= 15) whose attentional performance was measured daily (as described above) before, during and also 24-h after chronic access to nicotine self-administration under the differential dose procedure. Importantly, during chronic access to nicotine self-administration, attentional performance was measured immediately after each daily self-administration session to ensure that rats were still under the effects of nicotine.

This design uniquely allowed the monitoring of attentional performance at three different stages or states: before any exposure to nicotine self-administration (PRE), during the effects of self-administered nicotine (NIC) and 24-h after withdrawal (WD) from chronic nicotine self-administration under the differential dose procedure.

#### Experiment 3: effects of acute nicotine on nicotine withdrawal-induced attentional deficits

A subgroup of NoES (*n* = 11) and ES (*n*= 11) rats with a patent catheter from Experiment 2 was used in this experiment. Rats were tested for attentional performance 24-h after nicotine self-administration and 1 min after being administered intravenously a dose of nicotine. In total, animals received 3 doses of nicotine (0, 10, and 15 µg/injection, i.v.) using a within-subject counterbalanced Latin square design. Between each dose, animals were returned to nicotine self-administration under the differential dose procedure for 3 additional days before being withdrawn again from nicotine.

## Data Analysis

All data were subjected to relevant repeated measures ANOVAs, followed by Tukey post hoc tests where relevant. Statistical analyses were run using Statistica, version 7.1 (Statsoft Inc., Maisons-Alfort, France). Relationships between individual changes in attention and individual changes in nicotine intake escalation were assessed using Pearson correlations.

## Results

### Pre-existing individual variation in attentional performance predicts individual vulnerability to develop escalation of nicotine intake

We first determined attentional performance in animals trained and tested under a SD procedure (see Materials and Methods; Fig. 2). As previously shown [20], reducing the SD impaired attentional performance as revealed by both a decrease in accuracy (*F*_(2,80)_ = 29.1; *p* < 0.001; Fig. 2a), and an increase in omissions (*F*_(2,80)_ = 10.4; *p* < 0.001; Fig. 2a). Correct response latency also decreased with shorter SD of 0.5 s (*F*_(2,80)_ = 12.2; *p* < 0.001; Table 1), while incorrect response latency (*F*_(2,80)_ = 0.65; *NS*; Table 1) or latency to collect sucrose reward remained stable (*F*_(2,80)_ = 0.52; *NS*; Table 1), reflecting increased processing speed in this task rather than motor or motivational deficits. Moreover, an analysis of individual behavior revealed important individual variation, especially at the shorter SD of 0.5s, in accuracy (from 42.9% to 90%; variance = 146; median = 72.7%; Fig. 2b) and omissions (from 0% to 70.8%; variance = 407; median = 32.8%; Fig. 2c).

**Figure 2.**
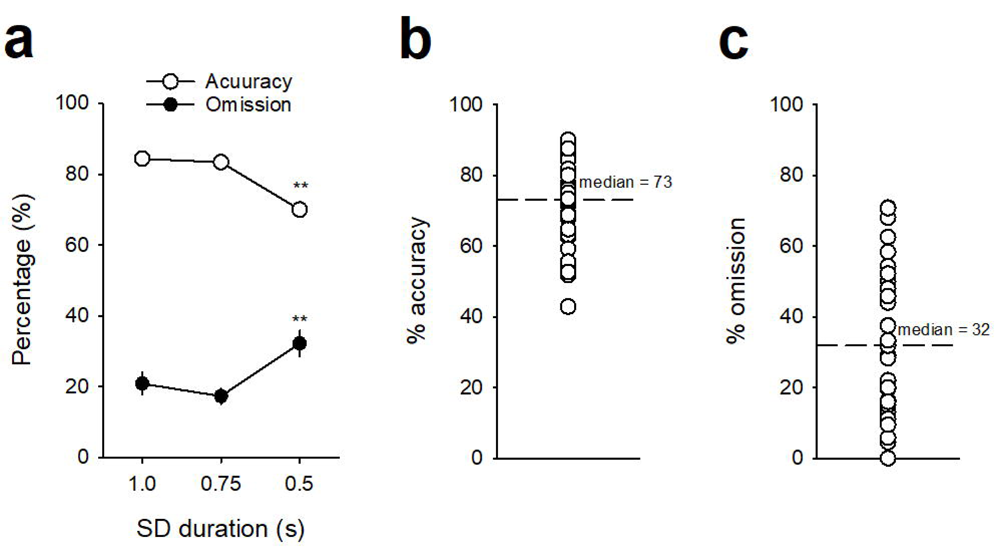
Attentional performance under the SD variable procedure. (**a**) Average percentage (mean ± SEM) of accuracy (*white circles*) and omission (*black circles*) under baseline condition as a function of the light stimulus duration, respectively SD1s, 0.75s and 0.5s. (**b**, **c**) Final individual percentage of accuracy (**e**) and omission (**f**) at the shortest SD0.5s. \*\**p* < 0.01, different from SD1s.

**Table 1:**
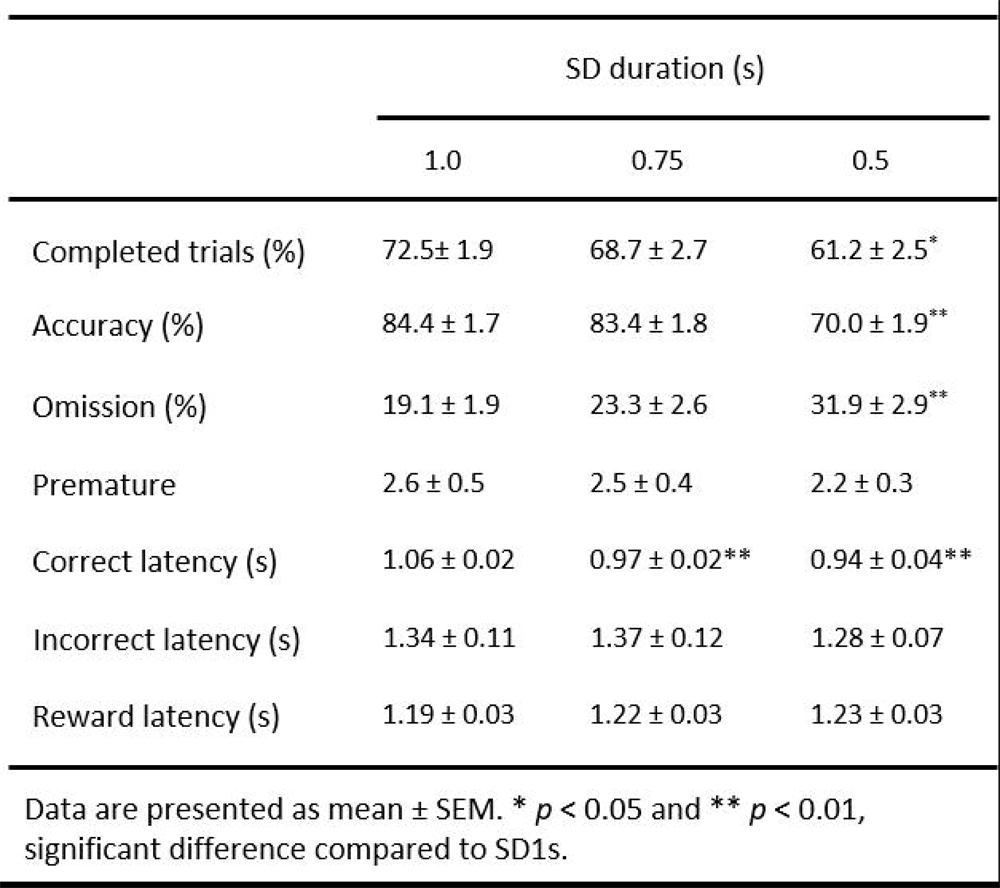
Attentional performance at SD variable.

We next assessed how individual variation in attentional performance can relate to individual vulnerability to nicotine intake escalation (see Materials and Methods for details; Experiment 1; Fig. 3). Animals were implanted with an intravenous catheter and subjected to a recently developed differential dose procedure of nicotine self-administration ([19]; see Materials and Methods for details). Consistent with previous findings, access to high doses of nicotine precipitated a rapid and robust escalation of nicotine intake in the escalation group (*F*_(2,78)_ = 18.1; *p* < 0.001; Fig. 3b). Though drug intake (i.e., average over the last 3 sessions at each dose) was similar in both groups at the initial dose of 30 µg/kg/injection (494 ± 69 µg/3h and 446 ± 46 µg/3h for NoES and ES groups, respectively; *NS*), nicotine intake first increased at 120 µg µg/kg/injection and further at 240 µg µg/kg/injection in the escalation group (from 446 ± 46 µg/3h to 1388 ± 105 µg/3h and 1867 ± 131 µg/3h at each dose; *p* < 0.001 at each dose), while it remained stable in the no-escalation group (from 494 ± 69 µg/3h to 632 ± 75 µg/3h and 598 ± 87 µg/3h at each dose; *NS*). At the end of the escalation procedure, the level of nicotine intake was 3 times higher in the escalation group than in the no-escalation group (598 ± 87 µg/3h and 1867 ± 131 µg/3h for NoES and ES groups, respectively; *p* < 0.001). Importantly, a correlation analysis across individuals revealed that individual changes in nicotine intake escalation were highly and positively correlated with individual scores in omissions (r = 0.67, *p* < 0.01; Fig. 3d) but not with accuracy (r = -0.21; *NS*; Fig. 3c). That is, animals with the highest scores of omissions escalated more their nicotine intake. This relationship was specific since latencies to respond (either correctly or incorrectly) or to collect the reward (Table 2), as well the number of visits and time spent in the drinking cup were not correlated with nicotine intake (r = -0.20 and r = -0.07; NS respectively). This suggests that differences in escalation level were not due to differences in motor ability or motivation.

**Figure 3.**
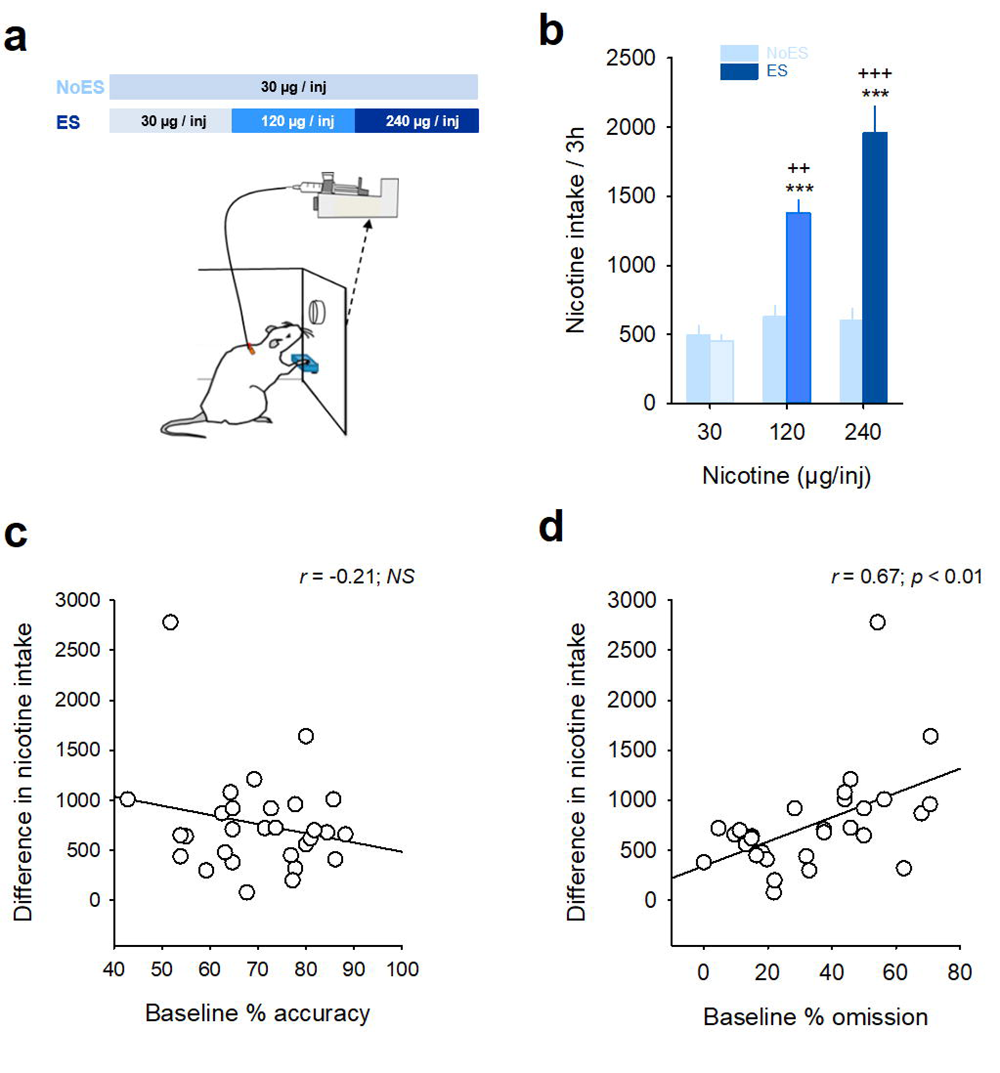
Pre-existing individual variation in attentional performance predicts individual vulnerability to develop escalation of nicotine intake. (**a**) Schematic representation and diagram of the differential dose procedure in the no-escalation group (NoES*, light blue*) that has access to the unit dose of 30 µg/kg/injection, and the escalation group (ES, *blue gradient*) that had access to increasing doses of nicotine (30 µg, 120 µg and 240 µg/kg/injection). (**b**) Average nicotine intake (mean ± SEM) during the last 3 days at each nicotine dose for the NoES (*light blue*) and the ES groups (*dark blue*). \*\*\**p* < 0.001, different from the lowest dose of 30 µg/injection. ^++^*p* < 0.01 and ^++^*p* < 0.001, different from the NoES group. (**c**, **d**) Individual changes in nicotine intake during escalation were highly correlated with individual % of omission (**d**) but not with individual % of accuracy (**c**).

**Table 2:**
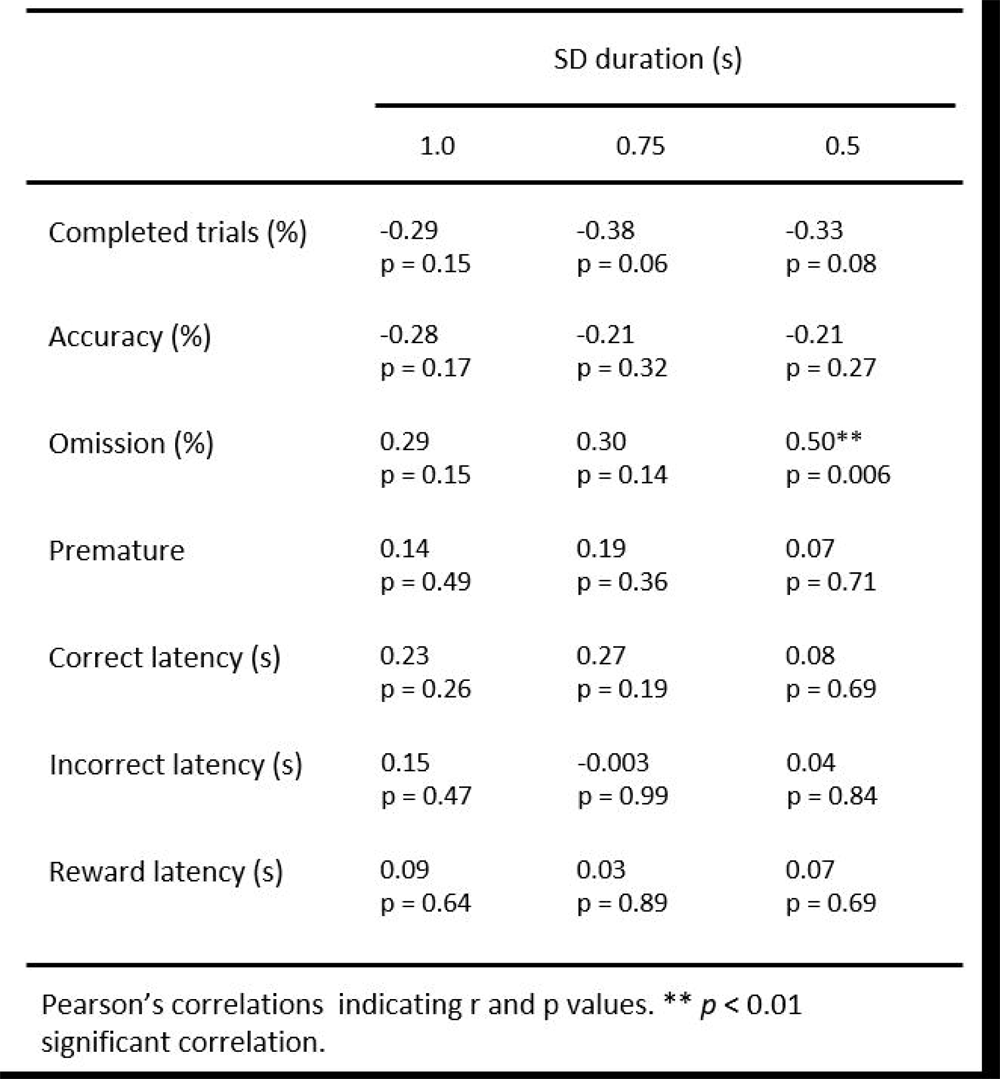
Correlation between 5-CSRTT performances and changes in nicotine intake during escalation.

|  | SD duration (s) |  |  |
| --- | --- | --- | --- |
|  | 1.0 | 0.75 | 0.5 |
| Completed trials (%) | -0.29<br>p = 0.15 | -0.38<br>p = 0.06 | -0.33<br>p = 0.08 |
| Accuracy (%) | -0.28<br>p = 0.17 | -0.21<br>p = 0.32 | -0.21<br>p = 0.27 |
| Omission (%) | 0.29<br>p = 0.15 | 0.30<br>p = 0.14 | 0.50**<br>p = 0.006 |
| Premature | 0.14<br>p = 0.49 | 0.19<br>p = 0.36 | 0.07<br>p = 0.71 |
| Correct latency (s) | 0.23<br>p = 0.26 | 0.27<br>p = 0.19 | 0.08<br>p = 0.69 |
| Incorrect latency (s) | 0.15<br>p = 0.47 | -0.003<br>p = 0.99 | 0.04<br>p = 0.84 |
| Reward latency (s) | 0.09<br>p = 0.64 | 0.03<br>p = 0.89 | 0.07<br>p = 0.69 |

Pearson's correlations indicating r and p values. \*\* $p < 0.01$ significant correlation.

### Attentional performance improves under the influence of self-administered nicotine but worsens during withdrawal

We next sought to determine whether nicotine self-administration and withdrawal can alter attentional performance. Another group of 27 animals (NoES, *n* = 12 and ES, *n*= 15) was trained under the SD and the differential dose of nicotine self-administration procedures as before (see Materials and Methods for details; Experiment 2; Fig. 4). Attentional performance was measured at three different stages or states: before any exposure to nicotine self-administration (PRE), immediately after daily nicotine self-administration sessions (NIC) and 24-h after withdrawal (WD) from chronic nicotine self-administration. Nicotine self-administration and withdrawal had little effect on accuracy (Fig. 4a) as nicotine withdrawal only decreased the percentage of accuracy compared to the self-administration period (day effect: *F*_(2,48)_ = 3.42; *p* < 0.05; PRE *vs* NIC, *NS*; PRE *vs* WD, *NS* and NIC *vs* WD, *p* < 0.05; Fig. 4a) in both groups (day x group: *F*_(2,48)_ = 0.33; *NS*) and for all SD (day x group x SD: *F*_(4,96)_ = 0.48; *NS*). In contrast, nicotine self-administration and withdrawal had a strong effect on omissions, especially in the escalation group (Fig. 4b). Nicotine self-administration decreased the percentage of omissions to a similar extent in both the no-escalation and the escalation groups (day effect: F_(2,48)_ = 17.9; *p* < 0.001 and day x group: *F*_(2,48)_ = 6.24; *p* < 0.01; PRE *vs* NIC, *p* < 0.05 for both NoES and ES groups) at all SD (day x group x SD: *F*_(4,96)_ = 0.48; *NS*), indicating that self-administered nicotine use increased attentional performance regardless of the dose available for self-administration. This improvement was also accompanied by a global decrease in the latencies to respond correctly (*F*_(2,48)_ = 7.8; *p* < 0.01; PRE *vs* NIC, *p* < 0.01; Table 3) and to collect sucrose reward (*F*_(2,48)_ = 27.9; *p* < 0.001; PRE *vs* NIC, *p* < 0.001; Table 3), but produced no change in premature responses (*F*_(2,48)_ = 2.0; *NS;* Table 3).

**Figure 4.**
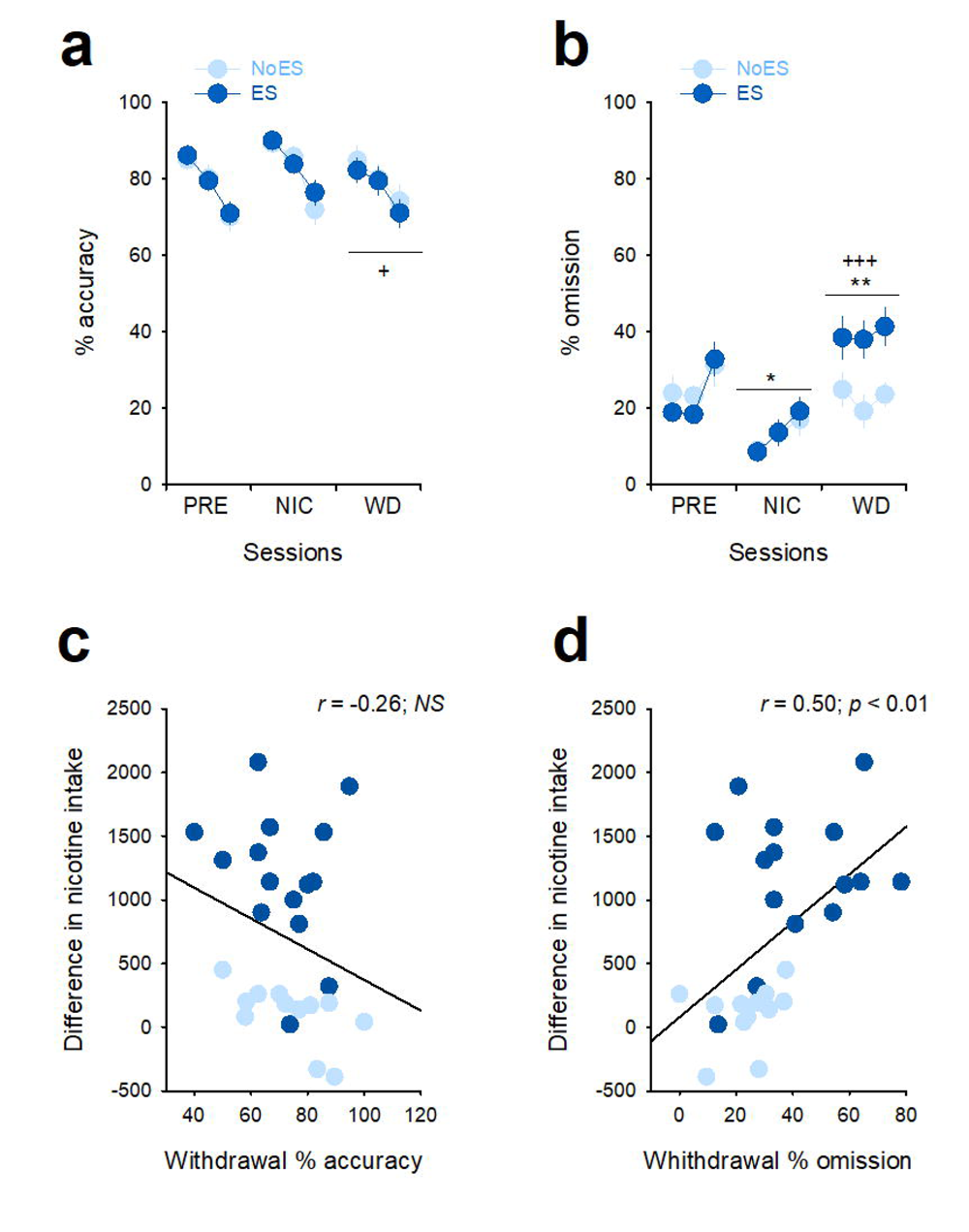
Attentional performance improves under the influence of self-administered nicotine but worsens during withdrawal. (**a**, **b**) Average percentage (mean ± SEM) of accuracy (**a**) and omission (**b**) before any exposure to nicotine self-administration (PRE), during the effects of self-administered nicotine (NIC) and 24-h after withdrawal (WD) from chronic nicotine self-administration in the NoES (*light blue*) and the ES groups (*dark blue*). For each session, the three circles represent the three light stimulus duration, respectively SD1s, 0.75s and 0.5s. \**p* < 0.05 and \*\**p* < 0.01, different from PRE. ^+^*p* < 0.05 and ^+++^*p* < 0.001, different from NIC. (**c**, **d**) Individual changes in nicotine intake were highly correlated with individual % of withdrawal induced-increase in omission (**d**) but not in accuracy (**c**).

**Table 3:**
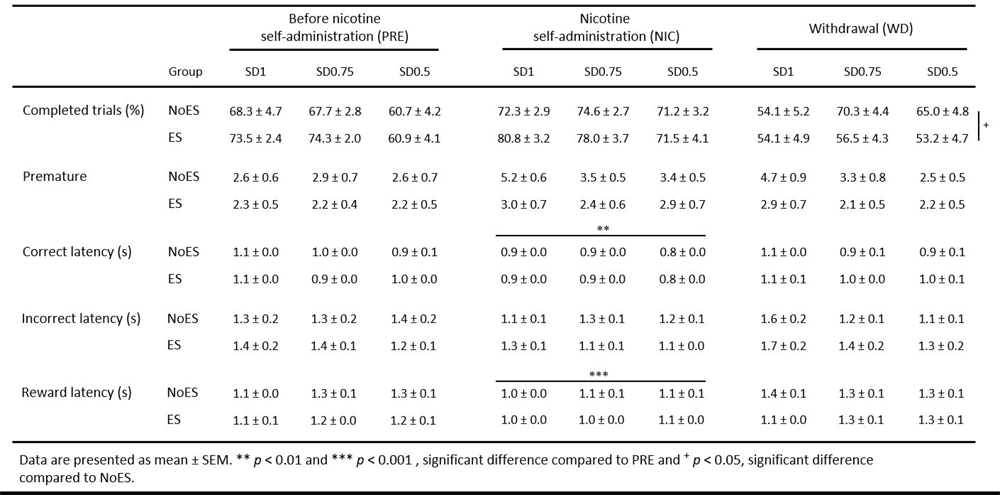
Changes in attentional performance after nicotine intake escalation (SA) and overnight withdrawal (WD)

In contrast, withdrawal from nicotine self-administration caused a large increase in omissions selectively in the escalation group (day x group: *F*_(2,48)_ = 6.24; *p* < 0.01; NoES-PRE *vs* NoES-WD, *NS*; NoES-NIC *vs* NoES-WD, *NS*; ES-PRE *vs* ES-WD, *p* < 0.01; ES-NIC *vs* ES-WD, *p* < 0.001 and NoES-WD *vs* ES-WD, *p* < 0.05), indicating that withdrawal from escalated levels of nicotine intake induced attentional deficits. This withdrawal-induced increase in omissions was remarkably specific since it was not associated with other behavioral changes, including: latencies of correct or incorrect responses (*F*_(2,48)_ = 7.8; *p* < 0.01; PRE *vs* WD, *NS* and *F*_(2,36)_ = 0.6; *NS,* for correct and incorrect latencies, respectively; Table 3), latency to collect sucrose reward (*F*_(2,48)_ = 27.9; *p* < 0.001; PRE *vs* WD, *NS*; Table 3), or premature responses (*F*_(2,48)_ = 2.0; *NS;* Table 3).

Moreover, individual changes in nicotine intake escalation were highly and positively correlated with changes in omissions during withdrawal specifically at the shorter SD (SD1s-WD: r = 0.22, *NS*; SD0.75s-WD: r = 0.43, *p* < 0.05; SD0.5s-WD: r = 0.50, *p* < 0.01; Fig. 4d), but not with changes in accuracy (SD1s-WD: r = -0.26; SD0.75s-WD: r = -0.16; SD0.5s-WD: r = -0.26; *NS* for all; Fig. 4c). Thus, the intensity of the attentional deficits induced by withdrawal from nicotine use was directly linked to the level of escalation of nicotine intake.

### Acute nicotine dose-dependently alleviates withdrawal-induced attentional deficits

Finally, we tested whether these withdrawal-induced attentional deficits can be alleviated by acute nicotine. A subgroup of animals from the previous experiment (*n* = 11 each for NoES and ES groups) received a passive intravenous injection of nicotine (0, 10 and 15 µg/injection, i.v.) 1 min before being tested for attentional performance 24-h after withdrawal from nicotine self-administration (see Materials and Methods for details; Experiment 3; Fig. 5). Acute administration of nicotine slightly increased accuracy at all SD during withdrawal in both groups at the higher dose used (dose effect: *F*_(2,40)_ = 3.25; *p* < 0.05; 0 *vs* 10 µg/inj, *NS* and 0 *vs* 15 µg/inj, *p* < 0.05 but dose x group: *F*_(2,40)_ = 2.62; *NS* and SD x dose x group: *F*_(4,80)_ = 0.64; *NS*; Fig. 4c). Most importantly, acute nicotine dose-dependently normalized the increase in omissions observed during withdrawal selectively in the escalation group (dose x group: *F*_(2,40)_ = 7.23; *p* < 0.01; ES-0 *vs* ES-10 µg/inj, *p* < 0.01 and ES-0 *vs* ES-15 µg/inj, *p* < 0.001; Fig. 4d), and at every SD (SD x dose x group: *F*_(4,80)_ = 0.59; NS), indicating that nicotine treatment can restore withdrawal-induced attentional deficits. This nicotine treatment also decreased latencies to respond correctly (*F*_(2,40)_ = 4.6; *p* < 0.05; 0 *vs* 10 µg/inj, *NS* and 0 *vs* 15 µg/inj, *p* < 0.05; Table 4) and to collect sucrose reward (*F*_(2,40)_ = 11.1; *p* < 0.001; 0 *vs* 10 µg/inj, *p* < 0.01 and 0 *vs* 15 µg/inj, *p* < 0.001; Table 4), without affecting the number of premature responses (*F*_(2,48)_ = 2.0; *NS;* Table 4). Thus, it is possible that during abstinence from escalated levels of nicotine intake, rats may seek to take nicotine in part to alleviate these cognitive deficits.

**Figure 5.**
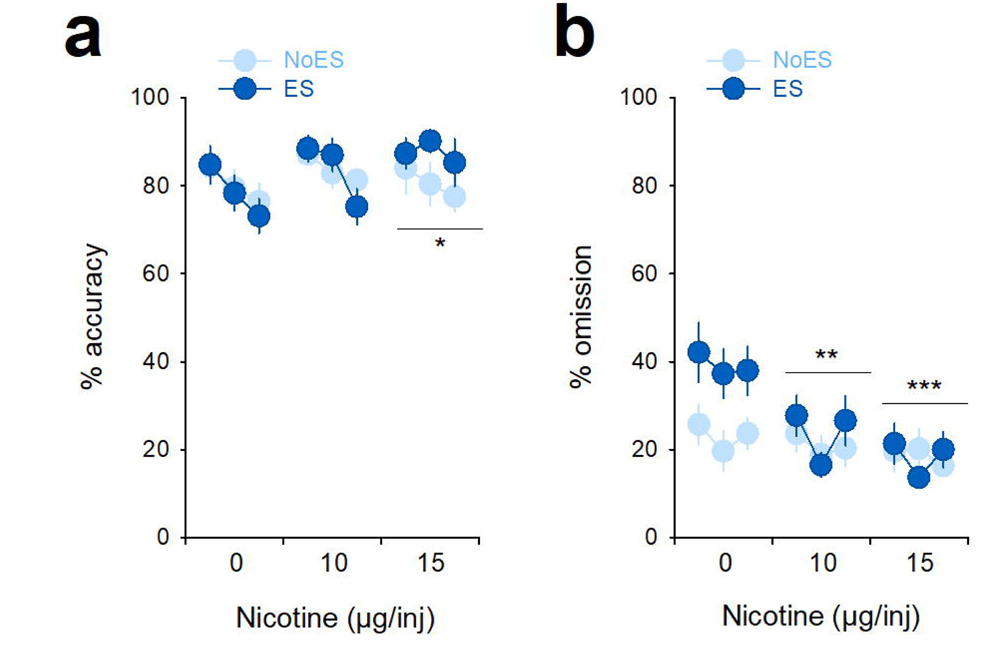
Acute nicotine dose-dependently alleviates withdrawal-induced attentional deficits. (**a**, **b**) Average percentage (mean ± SEM) of accuracy (**a**) and omission (**b**) after an acute passive intravenous injection of nicotine (0, 10 and 15 µg/injection, i.v.) administered 1 min before the SD procedure performed 24-h after withdrawal from self-administration. For each dose, the three circles represent the three light stimulus duration, respectively SD1s, 0.75s and 0.5s. \*\**p* < 0.01, different from 0 µg/injection.

## Discussion

Smoking addicts have deficits in frontocortical function and cognition even long after the discontinuation of drug use [3–6]. It is not clear, however, whether the cognitive deficits are a cause or a consequence, or both, of chronic nicotine use. Therefore, we characterized animals’ attentional performance both before, during and after escalation of nicotine intake using the 5-CSRTT.

We found that individual changes in nicotine intake escalation were highly and positively correlated with individual scores in omissions. As omissions reflect rats’ ignorance about location due to failure to pay attention to the stimulus presentation [20], this indicates that this pre-existing attentional feature predicts the propensity to escalate nicotine intake. This predictive relationship was specific since no other individual features in the 5-CSRTT were associated with escalation of nicotine intake in the present study. Notably, unlike some previous research, we did not find that individual differences in impulsive behavior in the 5-CSRTT (as measured by premature responses) can also predict individual variation in the rate of nicotine self-administration [21]. However, this apparent discrepancy is likely due to the specificity of our version of the 5-CSRTT which was not designed to study impulsivity. Indeed, in our study, attentional performance was assessed in rats that were well trained to wait and withhold their responses for a fixed and relatively long preparation time (i.e., 15 sec) and, thus, as a result, that made few premature responses.

We also found that the effects of self-administered nicotine improved attentional performance to the same extent regardless of the dose of nicotine available for self-administration. This outcome is relatively surprising since rats from the No-ES and ES groups maintain different levels of nicotine intake during self-administration. However, it is likely due to a saturation effect of nicotine on attentional performance [22]. This result extends previous evidence that nicotine administration via osmotic mini-pumps improved attention in the 5-CSRTT [1,13,14], and further supports the hypothesis that achieving this effect may contribute to tobacco smoking [2]. Moreover, and consistent with those studies, we found that self-administered nicotine reduced correct responses latency but had no effect on premature responses, indicating that an improvement in attentional processing and vigilance with no change in inhibitory control.

We also report that withdrawal from nicotine intake escalation induced attentional deficits. During 24-h withdrawal from nicotine self-administration, animals that had escalated their nicotine intake had a profound deficit in attentional performance, as indicated by a large increase in omissions. In contrast, animals that did not escalade their nicotine intake showed no deficits during withdrawal. Importantly, the selective increase in omissions seen during withdrawal from nicotine intake escalation could not be attributed to changes in non-cognitive modalities -such as sedation, locomotor impairment, or reduced motivation – since latencies to respond and to collect the reward were unaffected. Moreover, we found that the severity of attentional deficits during withdrawal depended on escalated levels of nicotine intake, worse attentional impairments being observed in animals that escalated their nicotine intake the most. Similar withdrawal-induced attentional deficits in the 5-CSRTT have been previously observed following extended, but not limited, access to self-administration of others drugs, including cocaine, amphetamine, and heroin [23–25], suggesting that common neurobiological substrates may underlie opiate-and stimulant-induced disturbances in visual attention. Finally, and consistent with behavioral studies in smokers [6,26], withdrawal-induced attentional deficits can be reversed by an acute dose of nicotine, supporting the deficit-compensating hypothesis of nicotine maintenance and resumption.

Overall, our findings indicate that while pre-existing individual difference in attentional performance predicts the vulnerability to escalate nicotine intake, part of the motivation for using nicotine during escalation might be to alleviate nicotine withdrawal-induced cognitive deficits, suggesting that both pre-existing and withdrawal-induced attentional deficits may contribute to initiation and maintenance of nicotine addiction.

## Author contributions

K.G. and S.H.A designed research and experiments; C.V.M. and K.G. performed behavioral experiments and associated data analysis; K.G. and S.H.A. wrote the paper.

